# Weighted gene co-expression network analysis reveals key genes and lncRNAs in desi cattle chronically infected with Johne’s disease

**DOI:** 10.1101/2025.01.17.633561

**Authors:** Abhisek Sahu, Mohd Abdullah, Saurabh Gupta, Shoor Vir Singh, Ankush Dhillon, Prabhati Yadav, Sarwar Azam

## Abstract

Johne’s disease (JD), caused by *Mycobacterium avium* subsp. *paratuberculosis* (MAP), is a chronic enteric disease with significant economic impacts on the dairy industry. To elucidate molecular mechanisms underlying MAP infection in *desi* cattle (*Bos indicus*), we conducted a comprehensive transcriptomic study. This revealed 1,905 protein-coding and 3,123 lncRNA genes as differentially expressed in MAP-infected samples, alongside the first comprehensive annotation of 45,947 lncRNAs in *desi* cattle. Weighted Gene Co-expression Network Analysis identified 11 co-expressed gene modules, with the turquoise module comprising 870 protein-coding genes and 934 lncRNAs, highly correlated with clinical traits. Functional enrichment analysis of this key module revealed significant involvement in defense response, inflammatory processes, and NK cell-mediated immunity and cytotoxicity. GSE analysis revealed suppressed pathways, including NK cell immunity and lectin response, facilitating bacterial persistence in chronic infection, while activated pathways included G-protein-coupled receptor signaling and metabolic pathways. Network approach identified 12 hub genes (**IL7R, TLR4, KLRK1, IFNG, TGFB1, CD68, CXCR6, GZMB, KLRG1, MMP9, GZMA, SELL**) associated with immune suppression, inflammation, and tissue remodeling. Additionally, 22 lncRNAs co-expressed with hub genes suggesting roles in modulating immune regulation. By identifying critical genes and lncRNAs, our study offers potential targets for developing innovative diagnostic and therapeutic strategies for JD.

## 1. Introduction

Paratuberculosis, also known as Johne’s disease (JD), is a chronic, insidious, and infectious disease characterized by incurable enteritis. It significantly affects the productivity of domestic ruminants, including cattle, while also infecting wild ruminants and a wide range of other animals, including primates and humans [1,2]. Symptoms of the disease are weakness, diarrhea, and a partial or complete loss of productivity [3,4]. The causative agent, *Mycobacterium avium* subsp. *paratuberculosis* (MAP), a robust, gram-positive, intracellular pathogen, exhibits remarkable resilience against diverse environmental and physical challenges [5,6]. It can endure adverse climatic conditions for extended durations outside the host and persists up to 55 weeks in dry and shaded environments [6]. Infected cattle experience a progressive disease course with four stages, leading to symptoms such as weight loss (with or without diarrhea), reduced milk and meat productivity, lethargy, hypoproteinemia, and emaciation [7–9]. These symptoms contribute to significant economic losses for dairy producers [10]. In India, the economic burden is exacerbated by the ban on cow slaughter in many states, rendering infected cattle unsalvageable. High morbidity rates and premature culling further amplify the losses [11,12]. Estimates from global studies highlight the financial impact, with annual losses per MAP-infected cattle ranging from $21 to $79 in the United States and approximately $40 in Canada [13]. Additionally, losses of $366 per lactating cow shedding MAP bacilli and up to $1,644 per 100 cows in herds with a 7% JD prevalence have been reported in the U.S., emphasizing the substantial economic implications of JD for dairy farming [14–16].

Research on MAP infection has focused on the transcriptomic responses of infected cattle to identify biomarkers and pathways involved in the disease. For instance, Marino et al. [17] utilized an in vitro model to investigate the early response to MAP infection using RNA-seq. Machugh et al. [18] analyzed gene expression in bovine monocyte-derived macrophages in response to in vitro MAP infection. David et al. [19] profiled gene expression in calves at 6 and 9 months post-inoculation with MAP, while Thirunavukkarasu et al. [20] identified novel pathways related to cholesterol and lipid metabolism in the early stages of MAP pathogenesis in cattle. Further studies by Shin et al. [21] and Hempel et al. [22] have examined immune responses, lipid metabolism, and the expression of immune regulatory genes in MAP-infected cattle. However, all of these studies have primarily focused on *Bos taurus* cattle. While these studies have provided valuable insights, their approaches have mainly employed differential gene expression analysis, potentially uncovering the host response and biomarkers for JD. However, they have not extensively explored co-expression modules of highly related genes, which are crucial for predicting gene function and identifying key genes involved in disease progression. Weighted Gene Co-expression Network Analysis (WGCNA) offers a robust method for integrating gene expression with phenotypic data, clustering genes with similar expression patterns into modules [23,24]. Genes within these modules often participate in related functions or pathways, making WGCNA a valuable tool for studying complex diseases [25]. This approach has been instrumental in identifying biomarker genes associated with various biological issues, including cancer [26,27], type I diabetes [28], rheumatoid arthritis [29], feed efficiency [30], and meat quality [31]. Its effectiveness in grouping genes into functional modules and uncovering regulatory mechanisms underlying complex traits has been emphasized in numerous recent studies.

India has a substantial cattle population of approximately 1.9 billion (https://dahd.nic.in/), primarily comprising a mix of breeds, with the majority being *Bos indicus*, also known as *desi* cattle [32]. *Desi* cattle are considered to possess better disease resilience compared to *Bos taurus*. However, JD is endemic among *desi* cattle, and high prevalence has been reported from many regions of the country [33–35]. Despite its substantial negative impact on dairy farming economics and potential public health implications, research on JD in *desi* cattle remains relatively limited. Previous transcriptomic studies in *Bos taurus* have provided valuable insights into host responses to MAP infection. However, a comprehensive understanding of the molecular mechanisms underlying MAP pathogenesis in *desi* cattle remains unexplored. This study leverages a weighted gene co-expression network analysis of transcriptomic data from chronically infected *desi* cattle to identify key regulatory genes and pathways involved in the host immune response to MAP infection. By integrating network analysis and functional enrichment analysis, we aim to uncover novel insights into the pathogenesis of MAP infection, which ultimately leads to JD. These findings may contribute to the development of targeted strategies for the prevention and control of MAP infection in *desi* cattle, mitigating its significant impact on animal health and the dairy industry.

## 2. Materials and methods

### 2.1. Phenotyping, Sample collection, and RNA sequencing

Careful selection of *desi* cattle was a critical aspect of this study to ensure accurate transcriptome profiling. Diseased and healthy female *desi* cattle were meticulously chosen based on specific exclusion and inclusion criteria as mentioned in **Additional file 1**. Each cattle recruited for the project underwent phenotyping for body condition score (BCS), clinical condition, and production parameters [36]. Samples were collected from phenotype cattle as per the guidelines of the Committee for the Purpose of Control and Supervision on Experiments on Animals (CPCSEA), India, with the Institutional Animal Ethics Committee (IAEC) approval. RNA sequencing methods are detailed in **Additional File 1**. Raw data were preprocessed using Fastp [37], processed with SortMeRNA [38], and resultant non-rRNA paired-end reads were retained as cleaned reads for further analysis.

### 2.2. Quantification and differential expression of mRNA

The *Bos indicus* reference genome (UOA_Brahman_1) and annotation (Ensembl v110) were used to extract protein-coding transcripts with GffRead [39]. Cleaned reads were aligned to these transcripts using Salmon [40] for transcript quantification, followed by gene-level quantification via the “tximport” R package [41]. Differential expression analysis was conducted using DESeq2 [42], with genes considered differentially expressed mRNAs (DEmRNAs) if p-adj < 0.05 and log2fc > 2 or < -2. PCA, heatmaps, MA, and volcano plots were generated using R packages to visualize gene expression patterns.

### 2.3. LncRNA analysis

For long non-coding RNA (lncRNA) identification, a customized LncEvo pipeline [43] based on Nextflow was used with raw reads and the annotated *Bos indicus* reference genome (UOA_Brahman_1). Cleaned reads were aligned to the reference genome using STAR [44], and transcript structures were assembled using Stringtie [45]. These were merged into a unified GTF file with Stringtie’s merge function and compared with the reference GTF using CuffCompare to exclude transcripts with error-prone class codes (c, e, p, s). Stringent filtering steps, detailed in **Additional File 1**, were then applied to refine the transcript set. The pipeline ultimately produced a curated set of potential lncRNAs in GTF and FASTA formats, ensuring reliable identification.

LncRNA abundance was quantified using Salmon, followed by data processing with the tximport R package to create a count matrix. LncRNAs with mapped read counts ≥10 were retained, and DESeq2 identified differentially expressed lncRNAs (DElncRNAs) based on p.adj < 0.05 and log2fc < −2 or > 2. PCA, MA, and volcano plots were used to visualize expression patterns.

### 2.4. Module trait analysis

The WGCNA R package [24] was used to identify significant gene modules and hub genes from DEmRNAs and DElncRNAs. Using VST transformed, normalized DESeq2 output and phenotypic trait data, a scale-free network was constructed by determining the optimal soft threshold power. Pearson correlation was used to create a similarity matrix, converted to an adjacency matrix, and then to topological overlap matrices (TOM). Gene modules were detected with the ’DynamicTreeCut’ function, and closely related modules were merged. Module Eigengenes (ME) were calculated, and their relationships with traits were visualized via heatmaps. Modules strongly correlated with traits were identified, and intramodular analysis pinpointed genes with high geneModuleMembership and geneTraitSignificance, offering insights into module-trait associations.

### 2.5. Functional enrichment analysis of key module

To identify biologically relevant Gene Ontology (GO) terms associated with MAP infection, key module genes were analyzed for GO enrichment using the clusterProfiler R package [46] with the *Bos indicus* database. A significance threshold of adjusted p-value < 0.05 was applied. The enriched GO terms were visualized and plotted using clusterProfiler package.

### 2.6. Identification of hub genes

Genes within the key module were identified based on high module membership values, measured by intramodular connectivity (KME or KIM) as described by Langfelder et al. [47]. Hub genes were selected with thresholds of ModuleMembership (|MM|) > 0.7 and geneSignificance (|GS|) > 0.3. A protein-protein interaction (PPI) network was constructed using the STRING database [48], retaining interactions with a confidence score ≥ 0.4. The cytoHubba plugin [49] in Cytoscape [50] ranked genes based on Degree, Maximum Clique Centrality (MCC), and Maximum Neighborhood Component (MNC), identifying the top 20 genes for each measure. Additionally, the MCODE plugin [51] identified densely connected regions (modules) with scores > 0.5. Hub genes common to both cytoHubba and MCODE analyses were considered the final set of hub genes.

### 2.7. Co-expression analysis of DEmRNA and DElncRNA in key module

To investigate co-expression between DElncRNAs and DEmRNAs, we identified cis-regulated DEmRNAs for each DElncRNA within the key module [52,53]. These cis-targets were defined as DEmRNAs located within a 100 kb window upstream or downstream of the DElncRNA [54] or overlapping with it, as identified using the ’bedtools window’ tool [55]. DElncRNAs co-expressed with their target DEmRNAs and present within the same key module (node weight > 0.05) were considered as co-expressed DElncRNAs and included in the network analysis.

## 3. Results

### 3.1. Phenotyping of control and clinically infected animals

In this study, animals were recruited based on specific inclusion and exclusion criteria outlined in the methods section. The phenotyping process involved assessing multiple criteria, including body condition score (BCS), clinical condition, and ZN (Ziehl-Neelsen) staining status for each animal. For the infected group, cattle underwent a phenotyping period of at least six months before selecting 20 cattle for blood sample extraction. These cattle consistently displayed positive ZN staining results, loose feces, and low BCS, confirming their infection status. Similarly, the healthy/control group was composed of animals that were phenotyped for more than 1 year before blood sample extraction. To be included in the control group, cattle had to consistently maintain good health and meet the criteria for being classified as healthy throughout the monitoring period. A total of 22 animals from the same gaushalas (cattle shelters) where infected animals were identified were used for blood sample extraction in the control group (**Table 1**). This careful selection and monitoring process ensured that the two groups, infected and healthy, were distinct and suitable for the experiment’s objectives.

**Table 1:** Sample phenotypes and clinical details included in this study.

| Sl. No | Sample Id | Breed name | Condition | BCS | ZN staining | Fecal |
| --- | --- | --- | --- | --- | --- | --- |
| 1 | dis_013R | Tharparkar | Diseased | 2 | 2+ | Loose |
| 2 | dis_04R | Tharparkar | Diseased | 1.5 | 3+ | Diarrhea |
| 3 | dis_07R | Tharparkar | Diseased | 1.5 | 2+ | Loose |
| 4 | dis_09R | Tharparkar | Diseased | 1.5 | 2+ | Loose |
| 5 | dis_17 | Tharparkar | Diseased | 2 | 3+ | Loose |
| 6 | dis_19 | Tharparkar | Diseased | 1.5 | 3+ | Soft |
| 7 | dis_26 | Hariana | Diseased | 2 | 1+ | Soft |
| 8 | dis_27 | Hariana | Diseased | 1.5 | 3+ | Loose |
| 9 | dis_29 | Hariana | Diseased | 3 | 2+ | Loose |
| 10 | dis_250 | Gir | Diseased | 2.5 | 2+ | Loose |
| 11 | dis_376 | Gir | Diseased | 3 | 2+ | Soft |
| 12 | dis_413 | Gir | Diseased | 2 | 2+ | Diarrhea |
| 13 | dis_438 | Gir | Diseased | 3 | 1+ | Soft |
| 14 | dis_446 | Gir | Diseased | 2.5 | 2+ | Loose |
| 15 | dis_231 | Sahiwal | Diseased | 1.5 | 2+ | Soft |
| 16 | dis_247 | Sahiwal | Diseased | 2.5 | 2+ | Soft |
| 17 | dis_402 | Sahiwal | Diseased | 3 | 2+ | Loose |
| 18 | dis_682 | Sahiwal | Diseased | 2.5 | 1+ | Soft |
| 19 | dis_825 | Sahiwal | Diseased | 2 | 2+ | Soft |
| 20 | dis_830 | Sahiwal | Diseased | 2.5 | 2+ | Soft |
| 21 | hel_2 | Tharparkar | Healthy | 3.5 | 0 | Normal |
| 22 | hel_4 | Tharparkar | Healthy | 3.5 | 0 | Normal |
| 23 | hel_5 | Tharparkar | Healthy | 3 | 0 | Normal |
| 24 | hel_6 | Tharparkar | Healthy | 3 | 0 | Normal |
| 25 | hel_7 | Tharparkar | Healthy | 3 | 0 | Normal |
| 26 | hel_9 | Tharparkar | Healthy | 4 | 0 | Normal |
| 27 | hel_11 | Hariana | Healthy | 3 | 0 | Normal |
| 28 | hel_27 | Haryana | Healthy | 3.5 | 0 | Normal |
| 29 | hel_31 | Haryana | Healthy | 4 | 0 | Normal |
| 30 | hel_39 | Haryana | Healthy | 3 | 0 | Normal |
| 31 | hel_47 | Haryana | Healthy | 3 | 0 | Normal |
| 32 | hel_383b | Gir | Healthy | 3.5 | 0 | Normal |
| 33 | hel_390 | Gir | Healthy | 3.5 | 0 | Normal |
| 34 | hel_444 | Gir | Healthy | 3 | 0 | Normal |
| 35 | hel_713 | Gir | Healthy | 3.5 | 0 | Normal |
| 36 | hel_887 | Gir | Healthy | 3 | 0 | Normal |
| 37 | hel_890 | Gir | Healthy | 3.5 | 0 | Normal |
| 38 | hel_451 | Sahiwal | Healthy | 3 | 0 | Normal |
| 39 | hel_481 | Sahiwal | Healthy | 4 | 0 | Normal |
| 40 | hel_605 | Sahiwal | Healthy | 3.5 | 0 | Normal |
| 41 | hel_804 | Sahiwal | Healthy | 3.5 | 0 | Normal |
| 42 | hel_808 | Sahiwal | Healthy | 3 | 0 | Normal |

**Note:** **BCS:** Body condition score; **ZN staining:** Ziehl-Neelsen staining

### 3.2. Genome wide identification of expressed transcripts in healthy and infected animals

Transcripts expressed in PBMCs from phenotyped animals were captured for the RNA-seq experiment. Sequencing was performed on 22 healthy and 20 diseased samples using illumina short reads. A total of 3,438.61 million reads were generated from these 42 samples and were initially processed through a quality filter, resulting in 3,225.84 million clean reads (**Table S1**). After the removal of ribosomal RNA, 2,790.64 million reads were obtained (**Table S2**). As a result, each sample had non-rRNA, high-quality cleaned data with a minimum of 24 million and a median of 64.18 million paired-end reads.

### 3.3. Identification of differentially expressed genes

To explore gene expression alterations in MAP-infected cattle, we analyzed the expression profiles of coding genes. RNA-seq data was mapped to mRNA transcripts **(Fig. S1A)**, resulting in the identification of 18,524 expressed genes. Differential expression analysis revealed 1,905 significantly altered genes (|log2FC| ≥ 2, adjusted p-value < 0.05) between diseased and healthy cattle. Hereafter these differentially expressed coding genes are referred to as DEmRNA. Among these, 1,588 DEmRNAs were upregulated and 317 were downregulated in the diseased condition. Principal component analysis (PCA) and hierarchical clustering demonstrated clear separation between diseased and healthy samples **(Fig. 1A, S1B)**. The diseased group exhibited greater variability compared to the healthy group, which clustered very tightly. Volcano and MA plots **(Fig. 1B, S2A)** visualized the extent of differential gene expression, highlighting the significant transcriptional response to MAP infection. A heatmap **(Fig. S2B**) further illustrated the expression patterns of the DEGs.

**Figure 1:**
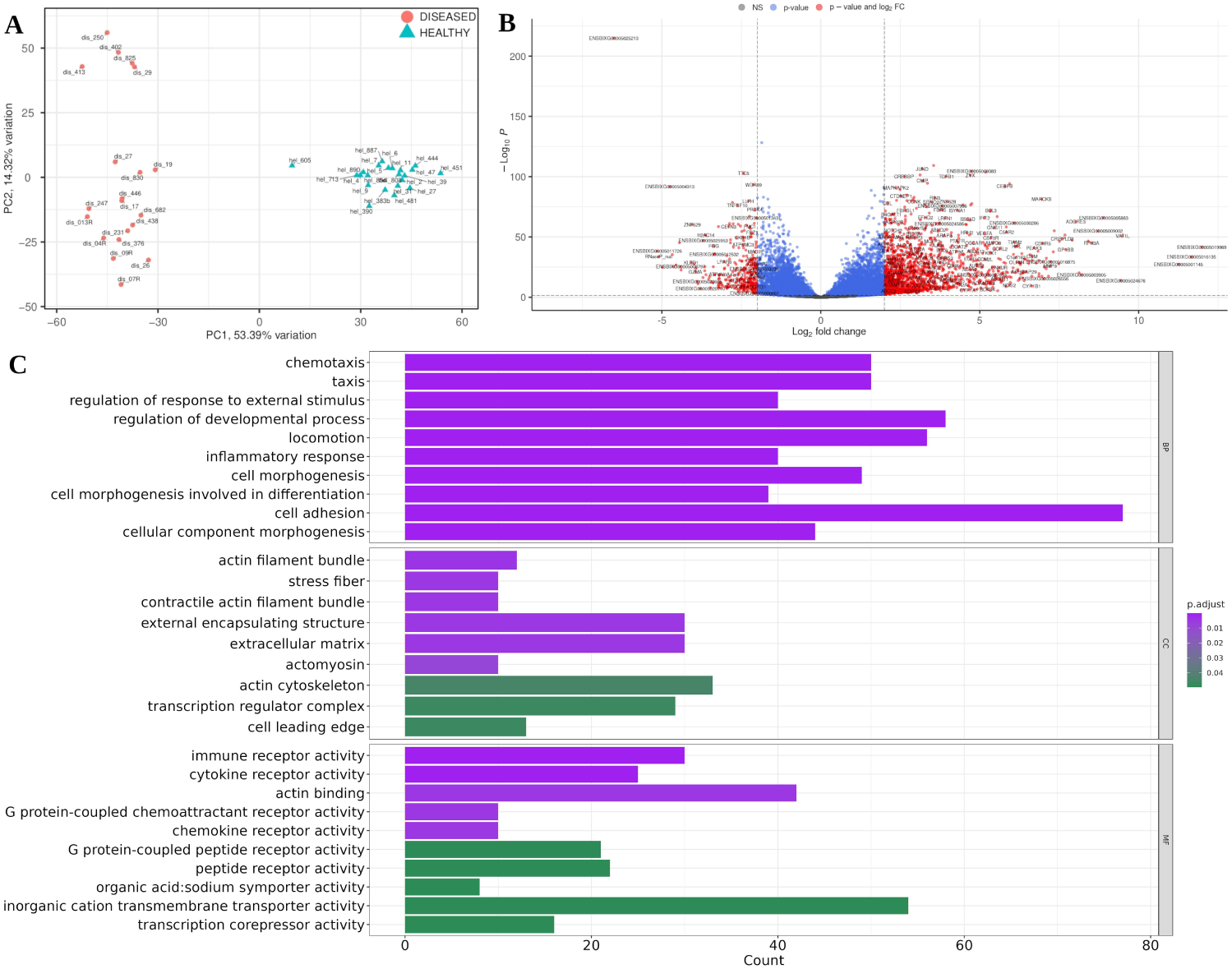
Clustering and differential gene expression analysis. **(A)** PCA plot of RNA-seq gene expression profiles. Triangles represent healthy individuals, while circles represent diseased individuals. **(B)** Volcano plot of differential gene expression analysis, with red points indicating significantly upregulated and downregulated genes based on log2 fold changes and p-value. **(C)** Bar plot of the top 10 enriched GO terms across categories: Biological Process (BP), Cellular Component (CC), and Molecular Function (MF).

A comprehensive GO analysis of 1905 DEmRNAs revealed insights into their biological processes (BP), cellular components (CC), and molecular functions (MF) **(Fig. 1C)**. Enriched BP terms were predominantly associated with immune responses, including inflammation, cell morphogenesis, and adhesion. Key MF terms were related to immune receptor and chemokine receptor activities. Gene Set Enrichment (GSE) analysis identified significant activation of signaling pathways, such as cGMP-mediated, cyclic nucleotide-mediated, and G protein-coupled receptor signaling pathways, and suppression of leukocyte-mediated cytotoxicity, lymphocyte-mediated immunity, natural killer cell mediated immunity and cell killing-related pathways in infected cattle.

### 3.4. Genome-wide identification and characterization of lncRNAs

A modified lncRNA prediction pipeline was employed to identify 58,621 lncRNA transcripts in cattle samples. Among these, 192 were known lncRNAs (class code "="), and the remaining were novel lncRNA candidates. These novel candidates were distributed in five classes based on their genomic location and overlap with annotated genes: intergenic (u), intronic (i), antisense (x), novel isoforms (j), and overlapping (o) **(Fig. 2A)**. The identified lncRNAs exhibited a wide range of lengths, from 200 bp to 22 kb. While many were multi-exonic, a significant number (11,042) were single-exonic. Among multi-exonic lncRNAs, the majority contained two exons (**Fig. 2B**). Only a small fraction exhibited a high exon count, with the maximum being 22 exons. The distribution of lncRNAs across chromosomes was uneven, with the X chromosome and chromosome 1 harboring the highest number of lncRNAs **(Fig. 2C)**.

**Figure 2:**
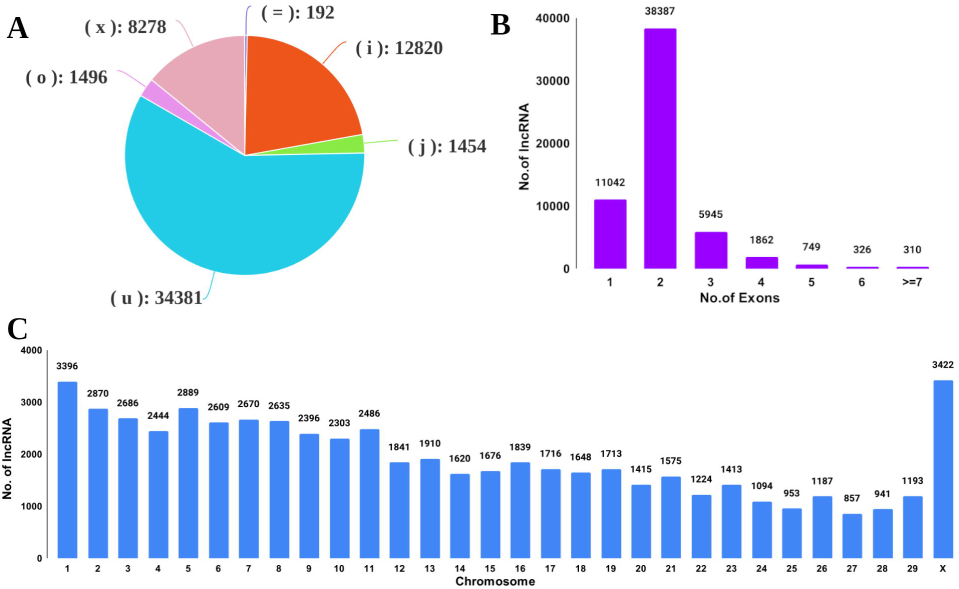
Genomic characteristics of lncRNAs in diseased vs. healthy *desi* cattle samples. **(A)** Distribution of predicted lncRNAs categorized by class code. **(B)** Exon count distribution among predicted lncRNA transcripts, illustrating the frequency of single and multi-exonic lncRNAs. **(C)** Chromosomal distribution of predicted lncRNA transcripts, showing the number of lncRNAs identified on each chromosome.

### 3.5. Identification of differentially expressed lncRNAs

For identification of differentially expressed lncRNAs, we quantified the expression levels of 58,621 lncRNA transcripts that corresponded to 45,947 genes using Salmon. After filtering for genes with a minimum read count threshold, 42,291 lncRNA genes were considered expressed. Differential expression analysis revealed 3,123 significantly altered lncRNA genes (|log2FC| ≥ 2, adjusted p-value < 0.05) between diseased and healthy cattle. These differentially expressed lncRNA genes will hereafter be denoted as DElncRNAs. Among these, 2,459 DElncRNAs were upregulated and 664 DElncRNAs were downregulated in the diseased condition. Volcano and MA plots (**Fig. S4A, S4B**) visualized the extent of DElncRNAs expression, highlighting the significant transcriptional response to MAP infection. A heatmap (**Fig. S4C**) further illustrated the expression patterns of the DElncRNAs, emphasizing the distinct clustering of diseased and healthy samples.

### 3.6. Construction of co-expression modules

A co-expression network was constructed using 1,905 DEmRNAs and 3,123 DElncRNAs with the WGCNA. After clustering all samples to identify outliers, no samples were excluded **(Fig. 3A**). A soft-threshold (β) power of 7 was chosen for a scale-free network (**Fig. 3B**), and genes were grouped into 18 modules through hierarchical clustering (**Table S3**). To refine module similarity, modules with a correlation >0.2 were merged, resulting in 11 final co-expression modules containing between 21 and 1,960 genes (**Table 2**, **Fig. 3C**). To investigate interactions within these co-expression modules, we generated a network heatmap illustrating their correlations, which indicated a relatively high degree of independence among clusters (**Fig. 4A**). To quantify the co-expression similarity across entire modules, we calculated their eigengenes and clustered them based on correlation. The resulting dendrogram showed that the 11 modules could be divided into two large subclusters, one comprising five modules and another with three modules, while two modules remained independent of either subcluster. This finding was further supported by the adjacency heatmap plot (**Fig. 4B**).

**Figure 3:**
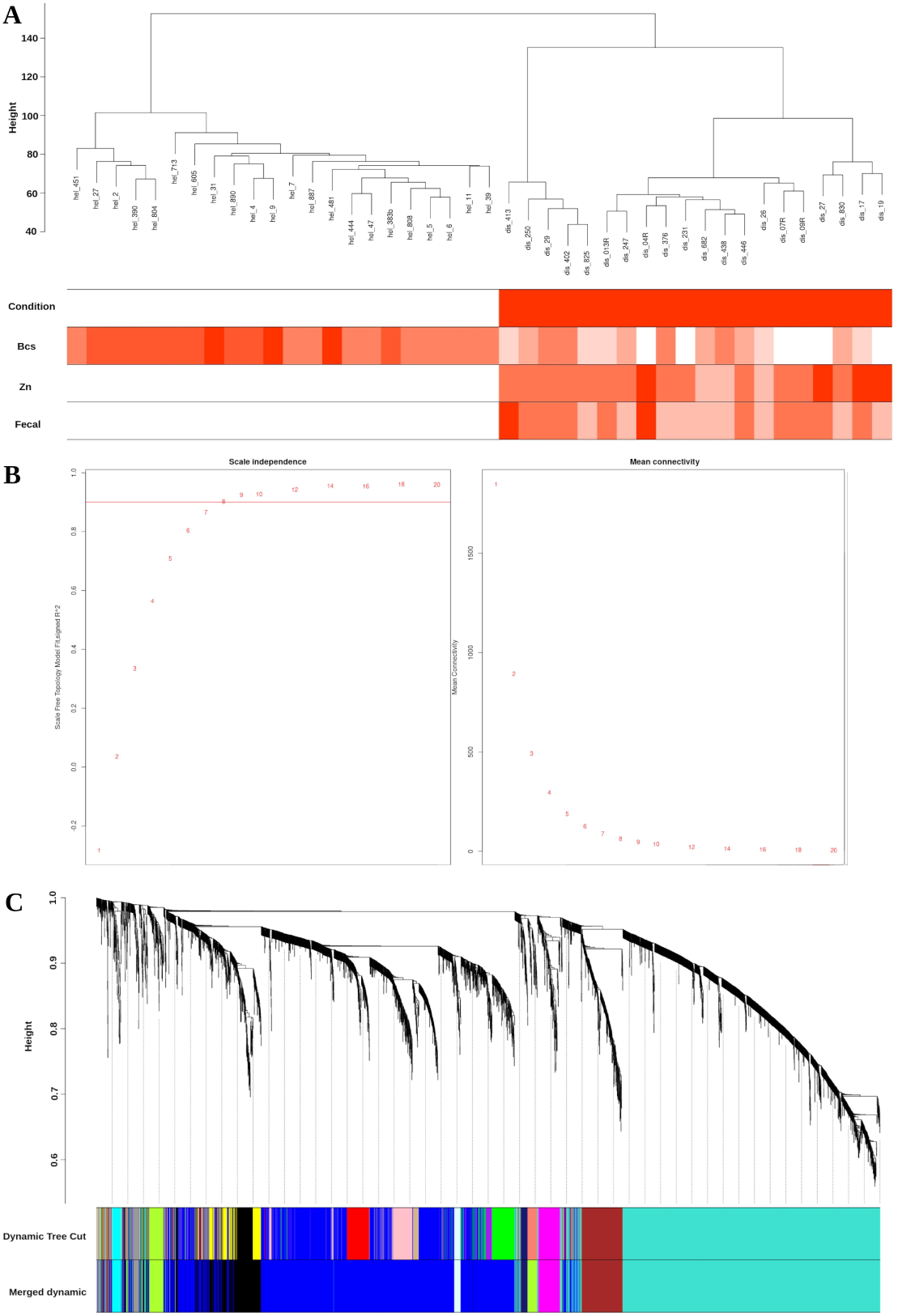
Construction of weighted gene co-expression network. **(A)** Sample dendrogram with an associated trait heatmap, where colors indicate the relative association with clinical traits. **(B)** Scale-free topology analysis, showing the fit index for various soft-thresholding powers (left) and mean connectivity for each soft-thresholding power (right). **(C)** Gene clustering dendrogram based on topological overlap, with colors representing distinct co- expression modules. Each module color indicates a specific gene cluster, with branches above representing individual genes. The top row displays the initial module assignment, while the bottom row shows the merged modules.

**Figure 4:**
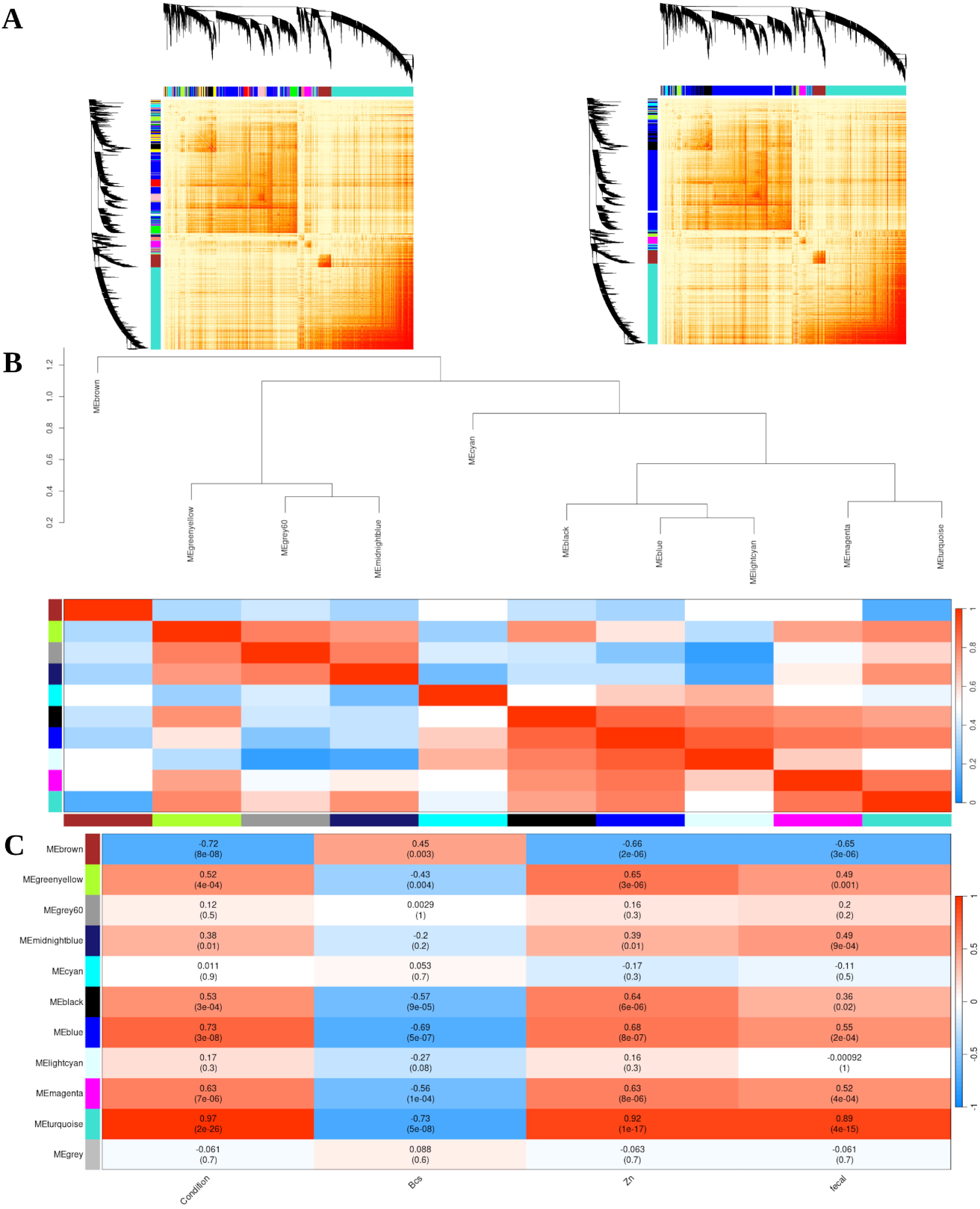
Module trait enrichment analysis.(A) Heatmap of the Topological Overlap Matrix (TOM) representing gene similarity in the weighted co-expression network. Lighter colors indicate lower overlap, while red indicates higher overlap. The left panel shows the TOM before module merging, and the right panel shows the TOM after merging modules. **(B)** Heatmap of eigengene adjacency, with color bars on the left and bottom indicating module assignments for each row and column. **(C)** Module-trait relationships, with each row representing a module eigengene and each column a trait. Each cell contains the correlation coefficient (R) and significance (p-value), with red indicating a positive correlation and blue indicating a negative correlation.

**Table 2:** Gene counts in co-expressed merged modules.

| Sl. No | Modules | Gene count |
| --- | --- | --- |
| 1 | Black | 434 |
| 2 | Blue | 1960 |
| 3 | Brown | 285 |
| 4 | Cyan | 60 |
| 5 | Greenyellow | 169 |
| 6 | Grey | 21 |
| 7 | Grey60 | 39 |
| 8 | Lightcyan | 56 |
| 9 | Magenta | 143 |
| 10 | Midnightblue | 57 |
| 11 | Turquoise | 1804 |

### 3.7. Identifying key modules

To identify key modules, we defined the representative expression profile for each module using the module eigengene. To explore relationships between these module eigengenes and specific traits of interest, we calculated correlation coefficients (**Fig. S5**). The resulting data were used to create a heatmap, visually depicting the correlation strength (R) and significance (p-value) between each module and the investigated traits (**Fig. 4C**). This heatmap highlights significant associations between modules and specific traits. Notably, the turquoise module, comprising 1,804 genes (870 protein-coding and 934 lncRNA), showed the most significant positive correlations with diseased conditions (**Fig. S6A**), fecal condition **(Fig. S6B)**, and ZN staining **(Fig. S6C)**, while displaying a negative correlation with BCS **(Fig. S6D)**. GS and MM analysis for the turquoise module (**Fig. S7**) confirmed the strong association between GS and MM. Based on its significant correlations with specific traits, the turquoise module was identified as the key module of interest for subsequent analysis.

### 3.8. Functional enrichment analysis of the turquoise module

GO enrichment analysis was performed on the 870 protein-coding genes in the turquoise module. The analysis revealed significant enrichment in BPs related to the regulation of immune responses, underscoring the module’s relevance to immune function. To provide a broader overview of BP enrichment, a word cloud was generated to highlight frequently enriched terms (**Fig. S8A**), while an induced GO enrichment graph demonstrated relationships among all enriched BP terms, revealing hierarchical patterns within immune response pathways (**Fig. S8B**). At the molecular level, genes within this module showed significant enrichment for processes such as defense response, inflammatory response, and natural killer cell-mediated immunity and cytotoxicity (**Fig. 5A**). To visually represent these enrichments, an enrichment map plot was created to organize the top 20 enriched BP terms into a network, with edges representing overlapping gene sets. The plot highlighted the interconnection of top enriched GO terms within the module (**Fig. 5B**). Additionally, a cnet plot was constructed to depict the distribution of specific genes contributing to key immune functions, offering a clear visualization of gene involvement (**Fig. 5C**). Several genes from the turquoise module, including TLR4 and IFT2, were enriched in all the top 10 GO terms.

**Figure 5:**
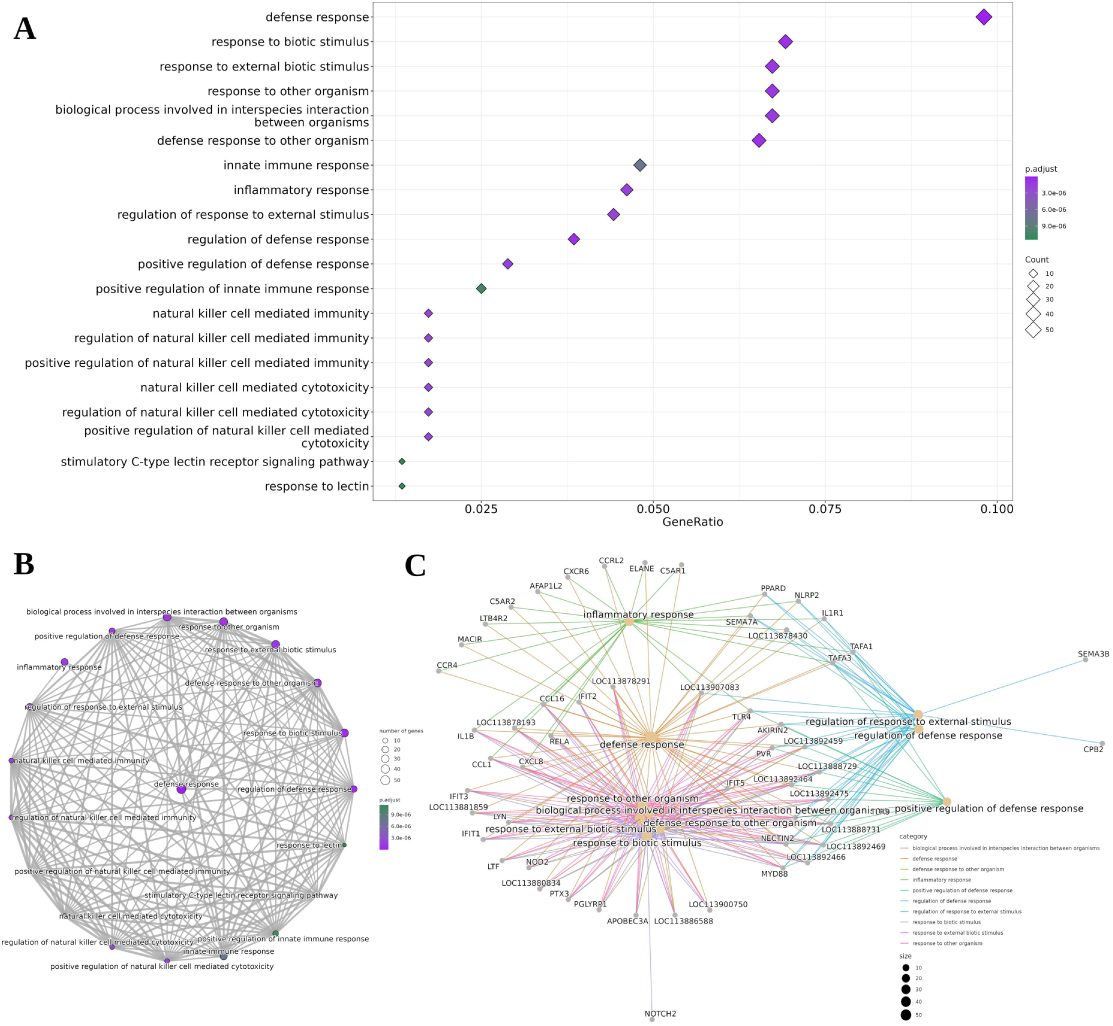
Functional enrichment analysis of the turquoise module. **(A)** Dot plot displaying the top 20 enriched GO BP terms associated with protein-coding genes in the turquoise module. **(B)** Enrichment map of the top 20 enriched BP terms, represented as a network, with edges indicating gene set overlap. **(C)** Category network plot (cnetplot) showing the relationships and overlap of genes across the top 10 GO biological process terms.

GSE analysis identified pathways significantly enriched in both upregulated and downregulated gene sets during infection. The top 30 enriched pathways are visualized using a GSE plot **(Fig. S9A)** and a ridge plot **(Fig. S9B)**. Suppressed pathways were predominantly associated with natural killer (NK) cell-mediated immunity, lectin response, nucleosome organization, and protein-DNA complex assembly. Conversely, activated pathways were linked to G-protein-coupled receptor signaling, circulatory system processes, metabolic processes, and other key biological functions.

### 3.9. Identification of hub genes and co-expressed lncRNA

To identify key regulatory (hub) genes in the turquoise module, a network-based approach was applied to the 870 protein-coding genes identified in the module. Filtering by MM and GS reduced the gene set to 526 genes. A high-confidence protein-protein interaction (PPI) network was then built using the STRING database configured for Bos indicus hybrids, resulting in 384 genes with interaction scores ≥0.4. The cytoHubba plugin ranked the top 20 most interconnected genes, while the MCODE plugin identified 34 genes from densely connected subnetworks. Combining these results, 12 hub genes (IL7R, SELL, TLR4, KLRK1, IFNG, TGFB1, CD68, CXCR6, GZMB, KLRG1, MMP9, and GZMA) were identified as central regulators in MAP infection, with high connectivity within the network **(Fig. 6A)**.

**Figure 6:**
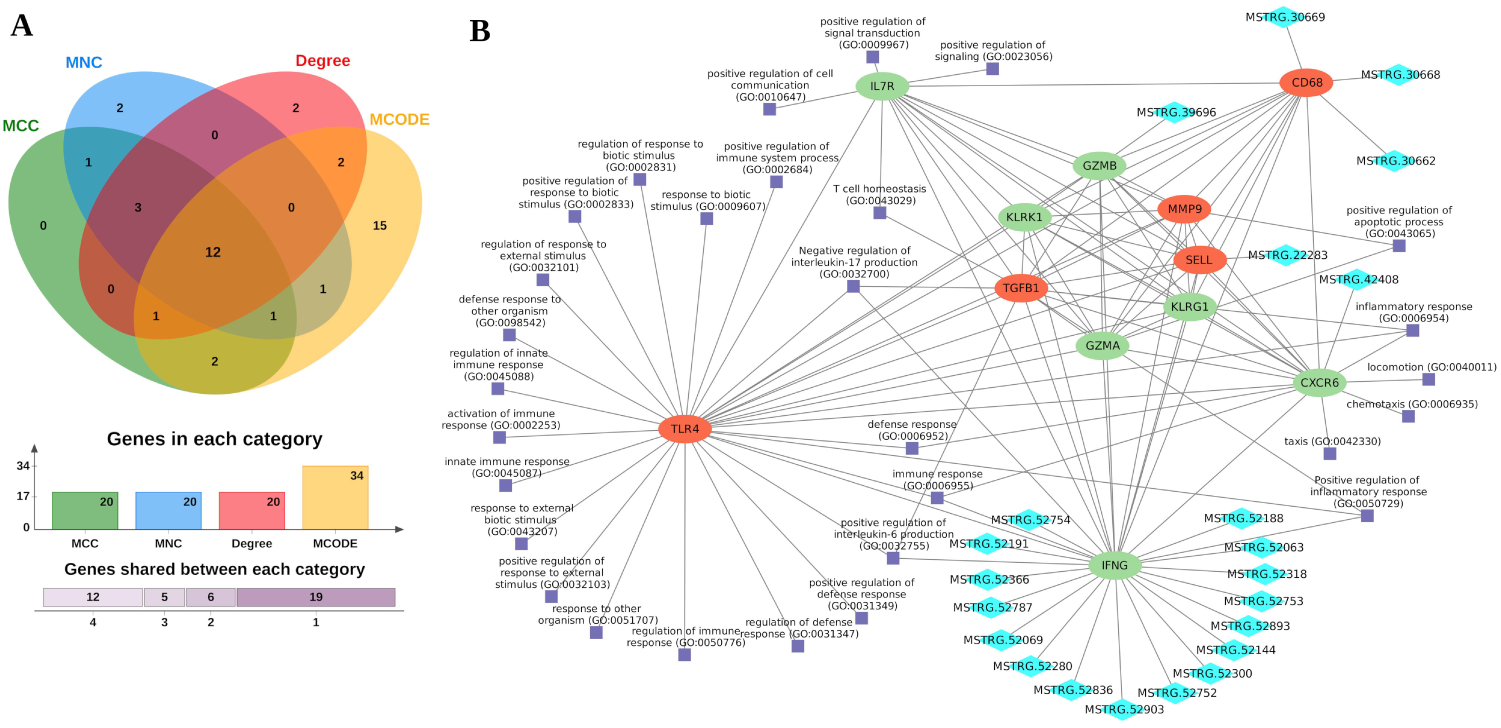
Identification of hub genes and their associated network of co-expressed lncRNAs and enriched GO terms. **(A)** Venn diagram illustrating the identification of a common set of hub genes across four algorithms. The overlapping region represents the hub genes shared by all four algorithms. **(B)** Network diagram depicting the interactions among hub genes, their co- expressed lncRNAs, and enriched GO biological process (BP) terms. Hub genes are shown as oval shapes, with upregulated genes highlighted in red and downregulated genes in green. Co- expressed lncRNAs are represented by diamond shapes in cyan, while enriched GO BP terms are displayed as square shapes in violet.

Recognizing the regulatory potential of lncRNAs, 934 DElncRNA genes from the turquoise module were investigated for their co-expression with hub genes. This analysis revealed 22 DElncRNAs co-expressed with five hub genes: IFNG (17 DElncRNAs), CD68 (3 DElncRNAs), CXCR6, GZMB, and SELL (1 DElncRNA each).These findings suggest that DElncRNAs may regulate hub gene expression, highlighting their potential roles in modulating immune response pathways during MAP infection (**Fig. 6B**).

## 4. Discussion

Previous transcriptomic studies on MAP infection in *Bos taurus* have primarily focused on identifying DEGs and candidate biomarkers [3,56,57]. While informative, these studies have often been limited in their ability to pinpoint core regulatory genes and pathways. This study advances our understanding of MAP infection in *Bos indicus*, a *desi* cattle known for its resilience. Leveraging a multi-layered approach, integrating phenotypic observations, transcriptomic data (mRNA and lncRNA), and WGCNA, we identified key regulatory hub genes and pathways associated with JD. These findings provide new insights into the complex host-pathogen interactions and offer a foundation for developing targeted therapeutic strategies.

The phenotypic characterization of the animals provided clear and clinically relevant distinctions between the chronically infected and control groups. The infected group exhibited consistent signs of MAP infection, including low BCS, loose feces, and positive ZN staining. This aligns with previous studies, which have identified these symptoms as hallmark indicators of clinical Johne’s disease [36]. RNA-seq analysis of infected and control groups revealed a robust transcriptional response to MAP infection in *desi* cattle, with 1,905 genes differentially expressed. Genes associated with enriched GO terms highlighted both activated and suppressed pathways in infected cattle. Activated signaling pathways included cGMP-mediated signaling, cyclic nucleotide-mediated signaling, and G protein-coupled receptor signaling. These pathways potentially assist the bacteria in persisting within the host by enhancing tolerance and antimicrobial resistance [58,59]. The cGMP signaling pathway, for instance, plays a crucial role in host survival against Gram-positive bacterial infections in *Drosophila* [60]. Elevated GPCR signaling has also been observed in *Mycobacterium tuberculosis* (Mtb) infections [61]. In contrast, suppressed pathways were primarily involved in adaptive and innate immune responses. Pathways such as leukocyte-mediated cytotoxicity, lymphocyte-mediated immunity, natural killer cell mediated immunity and cell-killing processes are essential components of cell-mediated immunity. The suppression of these pathways aids intracellular bacteria like MAP in evading key defense mechanisms that protect the host against intracellular pathogens [62]. Altogether, the altered transcriptomic response in chronically infected cattle suggests that MAP employs immune evasion strategies to persist within host cells, as previously reported [63].

LncRNAs are emerging as critical regulators of gene expression, particularly in response to diseases and stress [64]. However, identifying and functionally characterizing lncRNAs remains a significant challenge in livestock [65]. The current Ensembl reference genome annotation for *Bos indicus* lacks comprehensive lncRNA annotations. Previously, the only study by Sabara et al. [66] reported 10,360 lncRNAs in *desi* cattle using RNA-seq data from control and horn cancer samples. Our study provides a more comprehensive annotation by analyzing both MAP-infected diseased and healthy cattle. We identified 58,621 lncRNA transcripts in *Bos indicus* using 42 *desi* cattle. This number is comparable to the 47,683 lncRNA transcripts reported by Marete et al. [4] in *Bos taurus* in response to MAP infection. Further, differential expression analysis identified 3,123 DElncRNAs in MAP-infected cattle. Interestingly, a higher number of lncRNAs were upregulated in diseased conditions, suggesting a potential compensatory or regulatory mechanism activated during infection [67].

DEmRNAs and DElncRNAs exhibiting similar expression patterns were grouped into modules using co-expression network analysis. WGCNA, a widely employed co-expression network tool, has been successfully applied in numerous studies. A key advantage of WGCNA is its ability to assign an eigengene to each module, representing the overall expression profile. This eigengene can then be correlated with phenotypic data or clinical parameters, leading to highly reliable and biologically significant results. Using 1,905 DEmRNAs and 3,123 DElncRNAs, we discovered 11 modules of co-expressed genes. Associating these modules with specific traits requires accurate phenotypic data. An earlier study by Heidari et al. [56] on MAP infection in *Bos taurus* identified modules but lacked detailed phenotypic records, preventing module-trait analysis. In contrast, our study included clinical phenotype data, enabling us to identify the turquoise module as the key module most strongly associated with disease traits in *desi* cattle. This finding suggests that the genes within the turquoise module play a pivotal role in the pathological changes observed in MAP-infected *desi* cattle. The strong correlation between GS and MM for the turquoise module indicates that many of these genes are not only co-expressed but also highly relevant to the disease process.

Characterization of the turquoise module for biological significance revealed that 870 DEmRNAs were associated with top GO terms related to defense response and cell-mediated immunity, like other studies [56,68,69]. Additionally, GSE analysis highlighted the suppression of pathways involved in cell-mediated immunity. In contrast, activated pathways within the module included G-protein-coupled receptor signaling, circulatory system processes, and metabolic processes. The suppressed and activated pathways identified by GSE analysis from genes within the turquoise module were consistent with the enriched pathways identified using the overall set of DEmRNAs. This suggests that the turquoise module genes represent a core set of DEGs that effectively capture the key cellular changes associated with the disease. The observed activated pathways likely favor bacterial growth and persistence within the host, while the suppressed pathways represent the host’s defensive mechanisms that MAP attempts to evade.

We employed a network-based approach to identify key regulatory genes within the turquoise module, revealing 12 hub genes that emerged as central regulators of the network: **IL7R**, **TGFB1, IFNG**, **KLRK1**, **GZMB**, **GZMA**, **TLR4**, **CD68**, **MMP9**, **CXCR6**, **KLRG1**,

and **SELL**. These genes exhibited high |MM| > 0.7, |GS| > 0.3, and extensive protein-protein interactions (confidence score ≥ 0.4), underscoring their roles in the host-pathogen interplay during MAP infection. **IL7R** plays a pivotal role in immune suppression, with decreased expression linked to MAP susceptibility and inflammatory diseases like UC and IBD [4,70] . Its association with increased IFN-γ production upon MAP antigen stimulation further underscores its potential as a therapeutic target. Similarly, **TGFB1**, a pleiotropic cytokine, is essential for maintaining immune homeostasis. Its upregulation in MAP infection reflects a dual role: limiting excessive inflammation while contributing to immunosuppression, paralleling findings in TB-infected cattle [71,72]. **IFNG**, a critical cytokine for intracellular pathogen defense. It plays a pivotal role in activating macrophages and promoting a Th1-type immune response, which is essential for controlling mycobacterial infections [73,74]. As the disease progresses, a decline in IFN-γ levels is commonly observed, including in this study, resulting in a weakened immune response and increased susceptibility to opportunistic infections [75,76]. **KLRK1** (NKG2D), a receptor on NK cells and cytotoxic T cells, also exhibited downregulation, suggesting impaired cytotoxic immunity and favoring MAP persistence [77,78]. The serine proteases **GZMB** and **GZMA**, vital for immune-mediated pathogen clearance, displayed differential regulation [79] ; **GZMB** was downregulated, while **GZMA**, less studied in MAP, may have extracellular functions in immune modulation [80,81]. **TLR4**, an innate immune receptor, was upregulated, reinforcing its role in pathogen recognition and immune activation [82,83]. Genetic variations in TLR4 have been linked to MAP susceptibility, positioning it as a potential biomarker for diagnostic and therapeutic applications [84,85]. **CD68**, a macrophage marker, showed increased expression, indicative of macrophage activation and infiltration, though its association with granulomatous lesions reveals a complex role in disease progression [86–88]. **MMP9**, known for extracellular matrix remodeling and immune cell migration, was significantly upregulated, suggesting its utility as a transcriptional biomarker for JD [89,90]. Similarly, **CXCR6**, a G protein-coupled receptor, demonstrated context-dependent regulation, balancing immune cell recruitment with potential contributions to inflammation [91,92]. **KLRG1**, an immune checkpoint receptor associated with immune cell exhaustion, was identified as a novel hub gene, highlighting its likely involvement in MAP pathogenesis [93,94]. Lastly, **SELL** (L-selectin), crucial for leukocyte recruitment during inflammation [95], was identified as a hub gene, though its specific role in MAP infection warrants further exploration. Together, these findings highlight the intricate regulatory network underpinning MAP infection and offer potential targets for future research and therapeutic interventions.

Previous studies have emphasized the regulatory role of lncRNAs in gene expression, often mediated through their interactions with protein-coding genes. In bovine paratuberculosis induced by MAP, lncRNAs have been reported to influence the expression of adjacent genes related to immune response [56,96]. In our study, we identified 22 key lncRNAs that are differentially expressed and located near five hub genes, suggesting their role as molecular switches that fine-tune gene expression in response to MAP infection. These lncRNAs exhibit a high correlation with disease-associated traits, as indicated by their elevated GS scores, demonstrating their strong association with MAP infection-related traits. Among these, 17 lncRNAs are co-expressed with IFNG, a cytokine critical for macrophage activation and the promotion of Th1-type immune responses. This co-expression suggests that these lncRNAs may play a role in regulating IFN-γ expression, potentially influencing the host’s immune response to MAP infection. LncRNAs identified in the study could not be directly compared with those reported in previous studies on lncRNA involvement in MAP infection in cattle due to the absence of a comprehensive lncRNA database for cattle. This limitation represents a significant gap in the study, as such a resource would enable more robust comparisons, validation, and integration of findings across different studies.

## Conclusion

This study provides a detailed transcriptomic analysis of chronically MAP-infected *desi* cattle using RNA sequencing, identifying differentially expressed genes to explore the molecular mechanisms of MAP infection. WGCNA highlighted key gene modules, especially the turquoise module, strongly linked to clinical traits. GSE analysis revealed both activated and suppressed pathways in infected animals, reflecting the complex interaction between host and pathogen. The identification of hub genes such as IL7R, TLR4, KLRK1, IFNG, TGFB1, CD68, CXCR6, GZMB, KLRG1, MMP9, GZMA, and SELL provides insights into immune response regulation, immune suppression, inflammation, immune cell recruitment, and tissue remodeling in MAP infection. These genes may contribute to immune evasion and tissue damage during the disease process. Moreover, the study annotated a comprehensive set of lncRNAs in *desi* cattle and identified differentially expressed lncRNAs co-expressed with hub genes, suggesting their involvement in MAP pathogenesis. Overall, our findings suggest that MAP employs various strategies to evade host immune responses and establish persistent infection. The identified hub genes and lncRNAs represent potential biomarkers and therapeutic targets for the control of Johne’s disease. Further, validation and research into the exact mechanisms of these genes and lncRNAs in MAP pathogenesis are essential for advancing our understanding and treatment strategies.

## CRediT authorship contribution statement

**Abhisek Sahu:** Data curation, Formal analysis, Software, Visualization, Writing – original draft. **Mohd Abdullah:** Investigation, Resources, Validation. **Saurabh Gupta:** Project administration, Supervision. **Shoor Vir Singh:** Conceptualization, Project administration, Supervision, Funding acquisition, Writing – review and editing. **Ankush Dhillon:** Resources, Validation. **Prabhati Yadav:** Resources, Validation. **Sarwar Azam:** Conceptualization, Data curation, Formal analysis, Funding acquisition, Project administration, Supervision, Writing – original draft, Writing – review and editing.

## Acknowledgement

The authors gratefully acknowledge the financial support provided by the Department of Biotechnology (DBT), Ministry of Science, New Delhi, India. We also extend our sincere thanks to the National Institute of Animal Biotechnology (NIAB) for their invaluable support throughout this study. In particular, S.A. wishes to express gratitude to Dr. G. Taru Sharma, Director of NIAB, for her support. SVS is deeply indebted to the support of Dr. A.K.Gupta, Vice Chancellor and GLA university for this study.

## Data availability

The short-read sequencing dataset generated from the Illumina platform and used in this study has been submitted to the Indian Biological Data Centre (IBDC) under study id INRP000128. The detailed accession numbers for the RNA-seq data are provided in **Table S4**. All the data can also be accessed from the NCBI.

## Additional Files

**Additional File 1:** Additional methods

---

**Figure S1:**
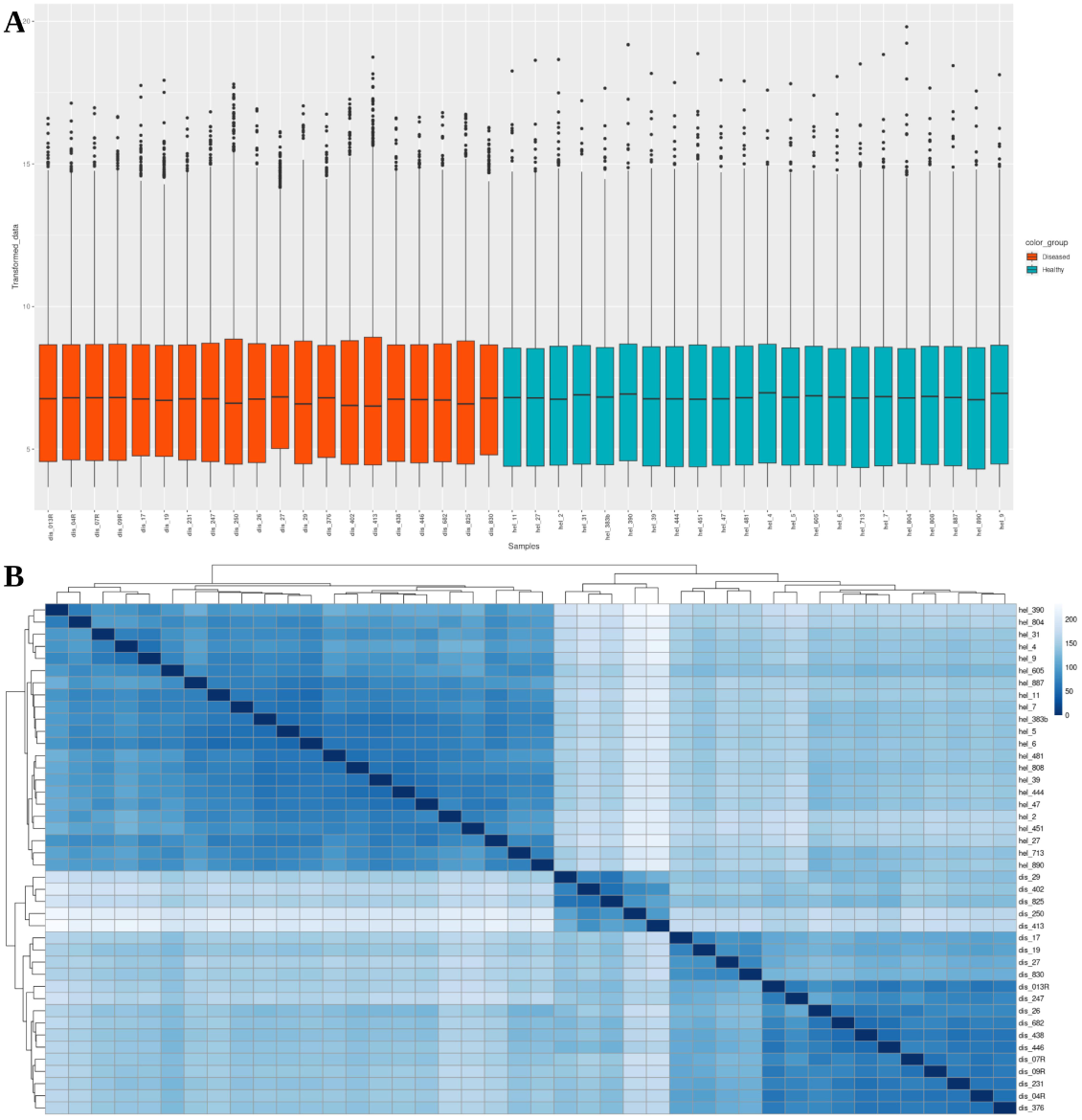
Data normalization and sample clustering. **(A)** Boxplot of transformed and normalized counts for each sample, illustrating consistent distribution across samples. **(B)** Hierarchical clustering dendrogram of samples based on gene expression patterns, showing distinct clusters that separate diseased and healthy samples.

**Figure S2:**
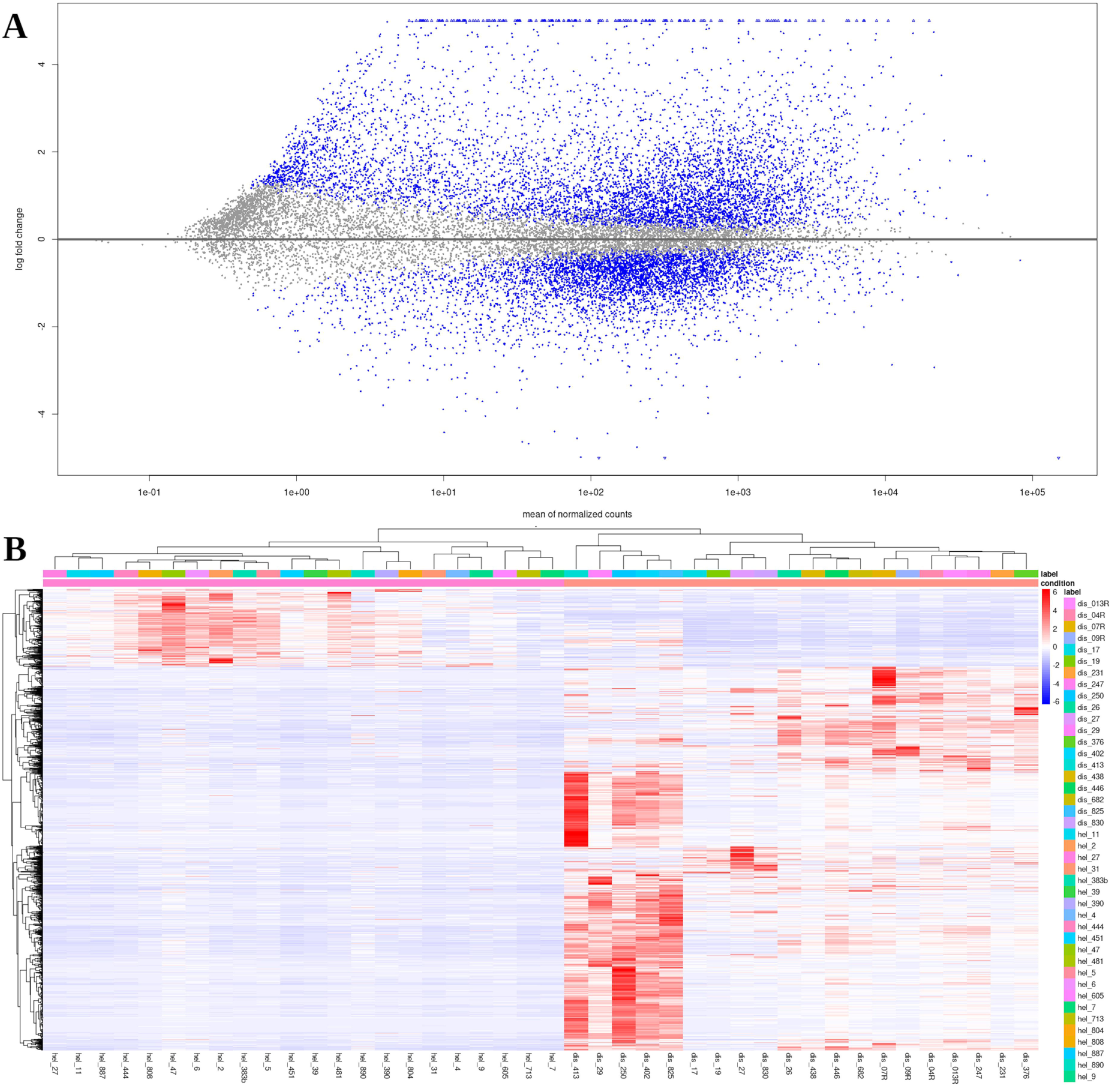
Differential expression of mRNA. **(A)** MA plot showing the relationship between the average expression (log-transformed mean counts) and the log2 fold change of differentially expressed mRNAs between disease and healthy conditions. **(B)** Heatmap displaying the expression patterns of differentially expressed genes across disease and healthy samples. Each row represents a gene, and each column represents a sample.

**Figure S3:**
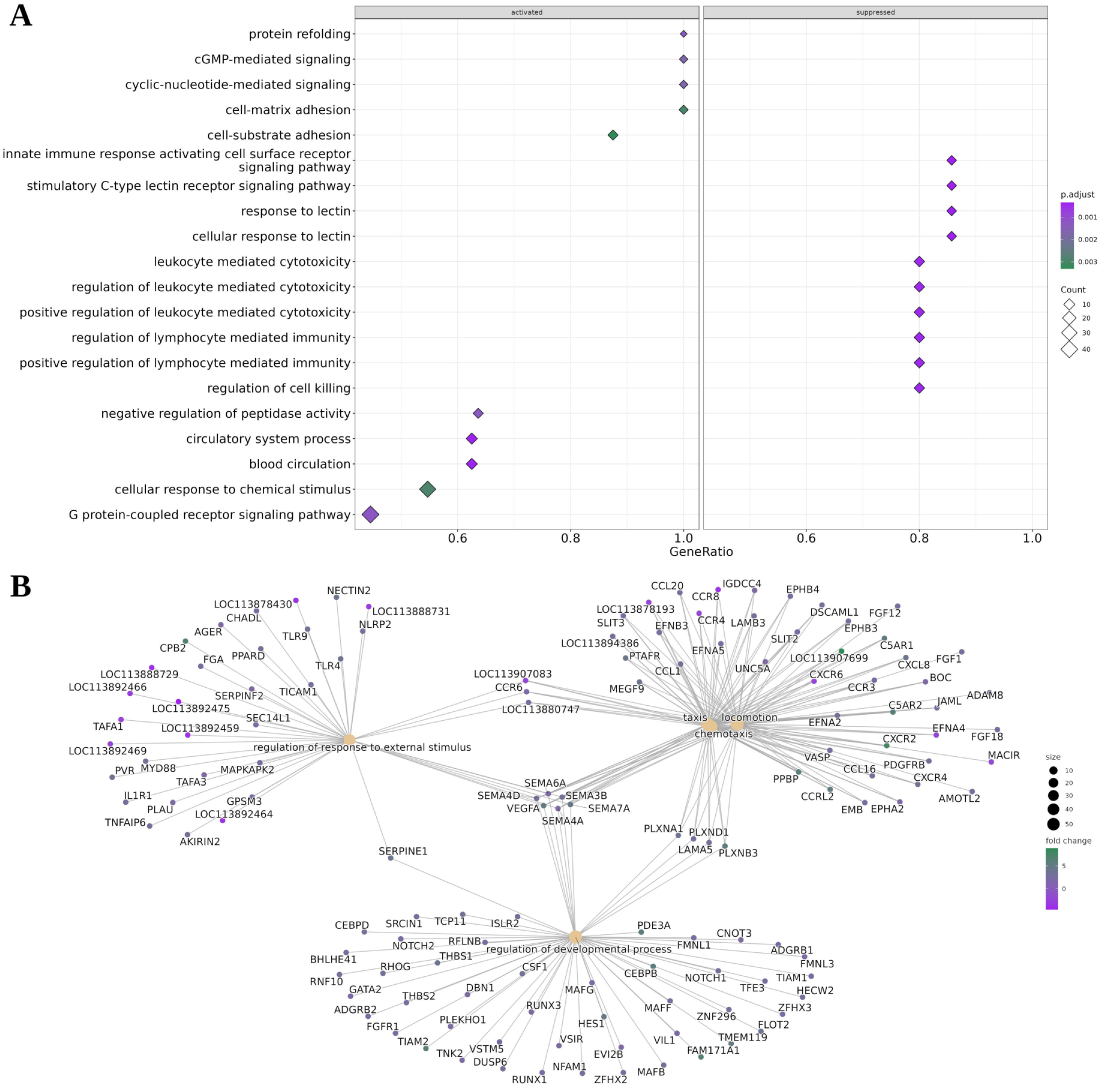
Enrichment of differentially expressed genes. **(A)** GSE plot displaying the top 20 pathways activated or suppressed in response to MAP infection. **(B)** Cnet plot showing the top five enriched GO terms and the genes associated with BP.

**Figure S4:**
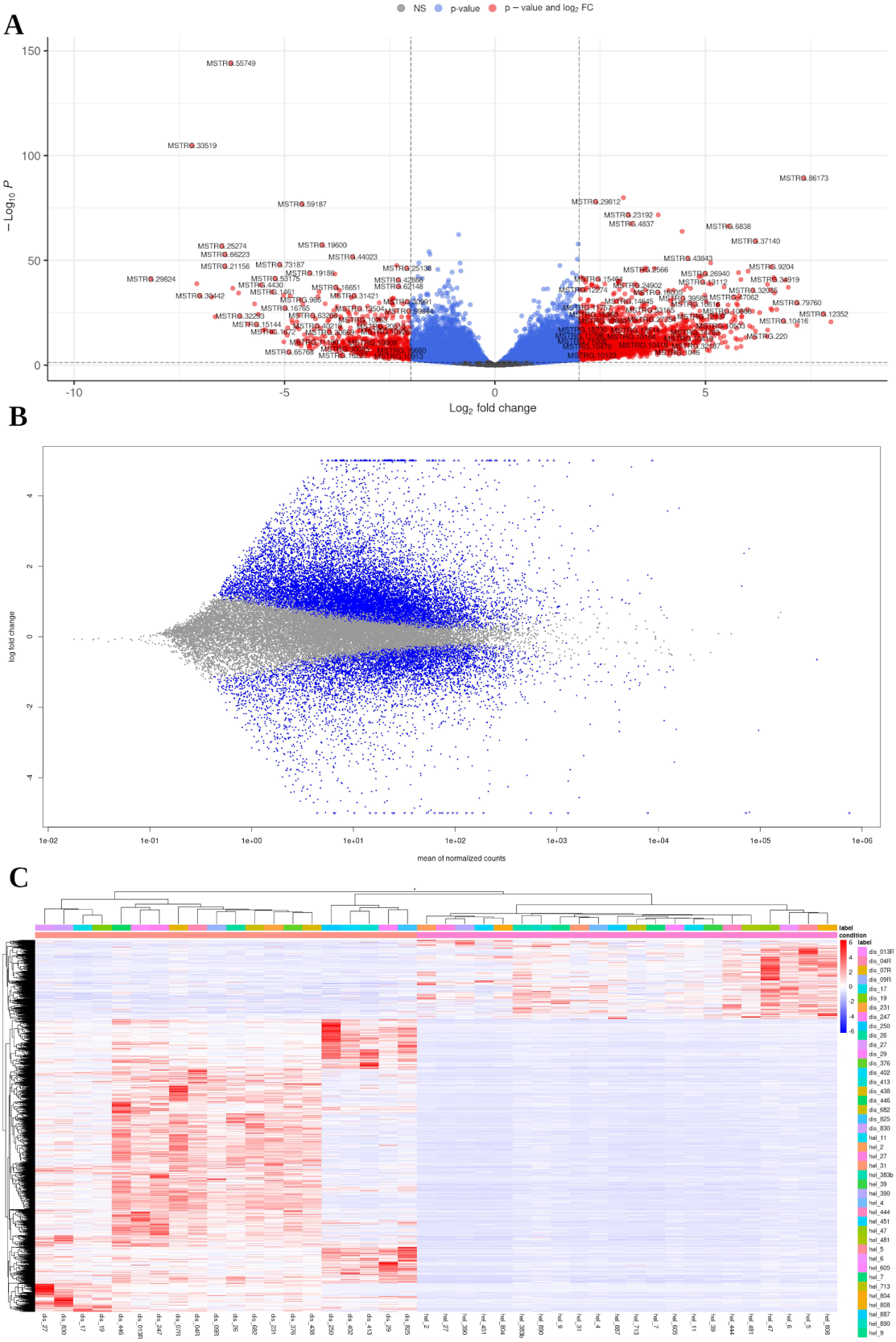
Differential expression of lncRNAs. **(A)** Volcano plot showing differentially expressed lncRNAs, with red points indicating upregulated and downregulated genes based on the log2 fold change (log2(FC)) between disease and healthy samples. **(B)** MA plot displaying the relationship between average expression (log-transformed mean counts) and the log2 fold change of differentially expressed lncRNAs between disease and healthy conditions. **(C)** Heatmap of differentially expressed lncRNAs in healthy versus diseased samples.

**Figure S5:**
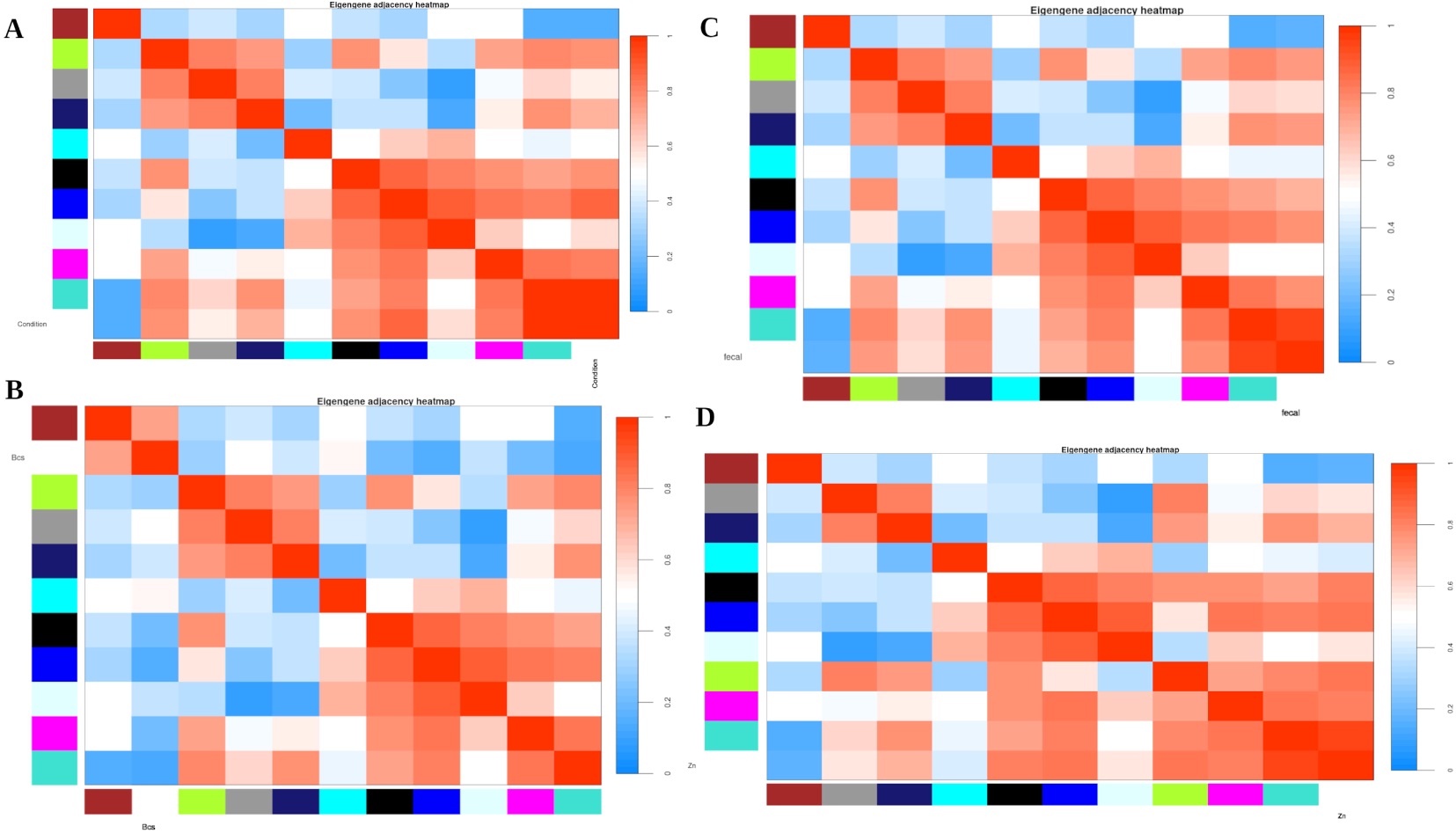
Eigengene heatmap showing correlated modules with traits. Heatmap of module eigengenes to identify clusters of correlated eigengenes (modules) in relation to each trait. **(A)** Condition **(B)** BCS **(C)** Fecal **(D)** ZN staining

**Figure S6:**
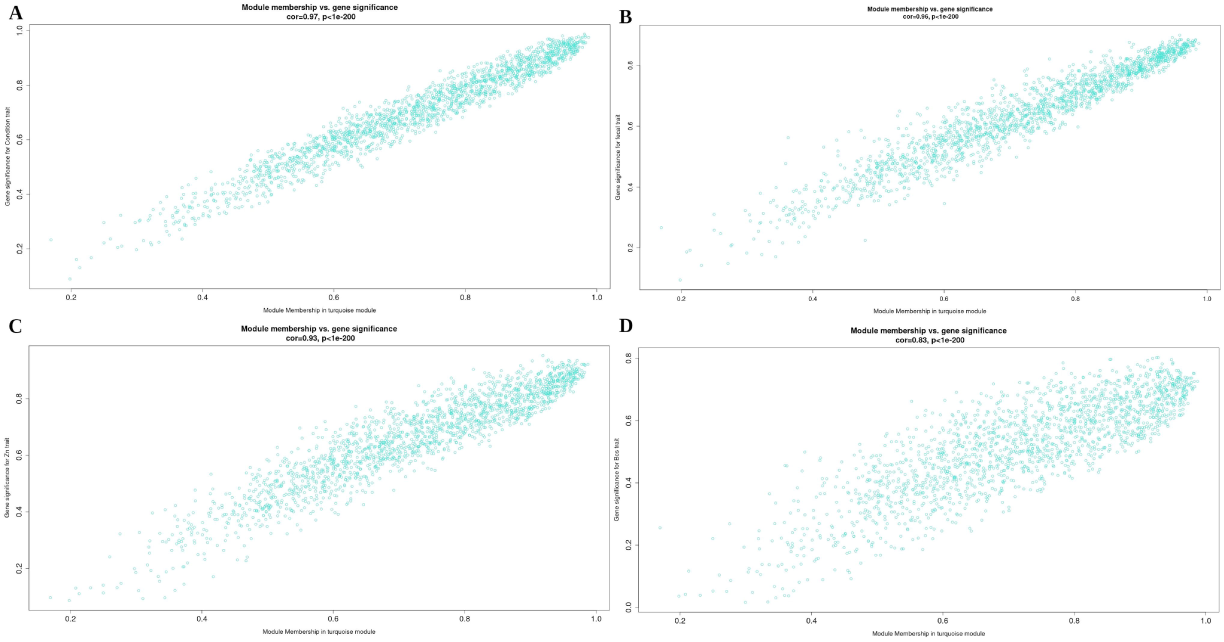
Heatmap of MM and GS in the turquoise module. **(A)** MM vs. GS with condition trait **(B)** MM vs. GS with fecal trait **(C)** MM vs. GS with ZN staining trait **(D)** MM vs. GS with BCS trait

**Figure S7:**
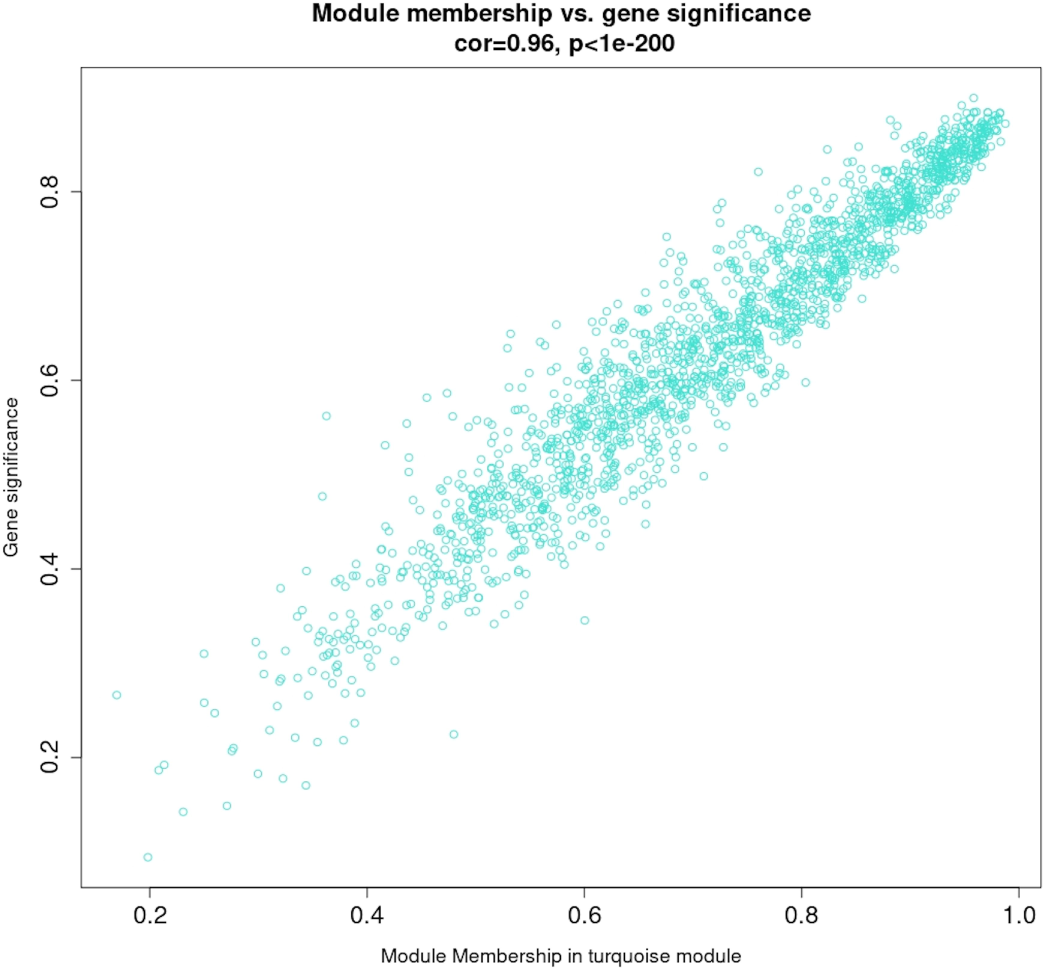
Relationship between MM and GS for MAP infection in turquoise module. Scatter plot showing the correlation between MM and GS in the turquoise module for MAP infection, comparing diseased and healthy cattle.

**Figure S8:**
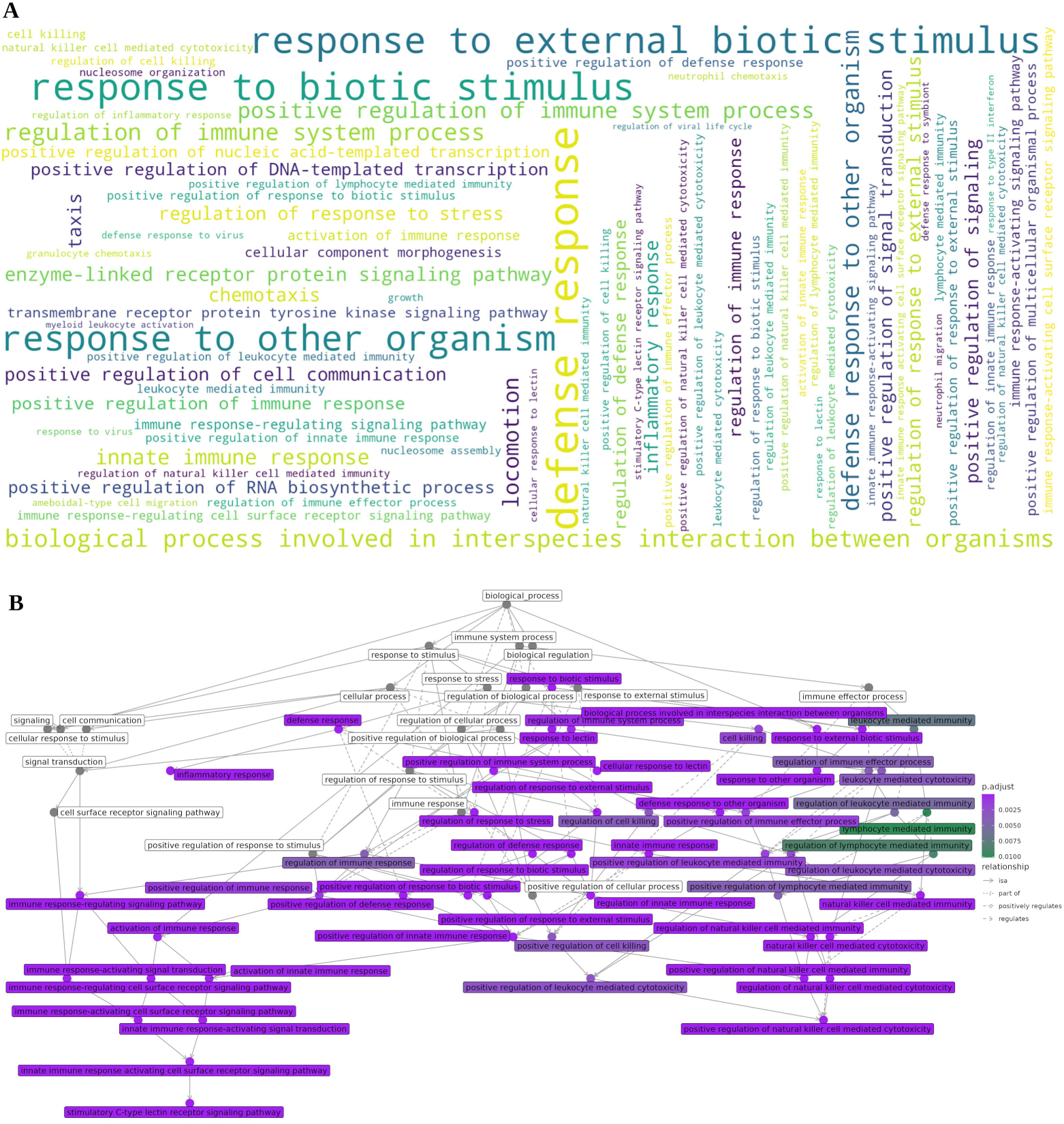
GO enrichment visualization of protein coding genes in turquoise module. **(A)** Word cloud of all enriched GO BP terms, with font size representing the frequency and significance of biological processes associated with protein coding genes. Larger terms indicate higher enrichment significance. **(B)** Enriched GO network graph showing the relationships among the top GO terms in BP. Nodes represent enriched GO terms, with edges indicating functional relationships based on shared genes. Node size reflects significance, and color intensity represents gene count. This graph highlights interconnected pathways.

**Figure S9:**
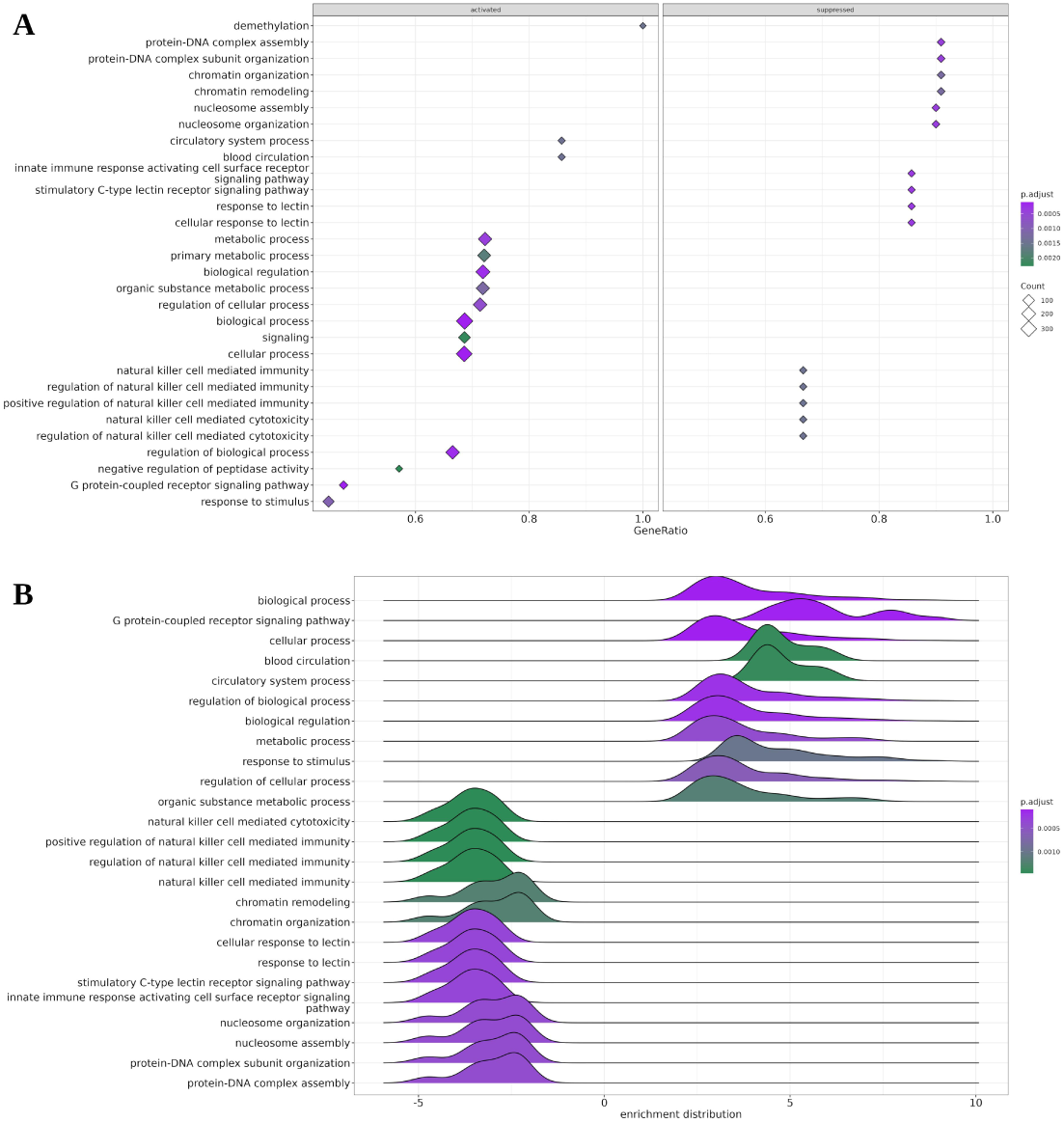
Gene set enrichment analysis. **(A)** GSE plot illustrating the top 30 enriched pathways, highlighting those that are activated or suppressed in the dataset. **(B)** Ridge plot showing the distribution of the top 25 upregulated and downregulated pathways, providing insight into the variability and significance of pathway changes.

