## Additional file 1 for "Weighted gene co-expression network analysis reveals key genes and lncRNAs in desi cattle chronically infected with Johne’s disease"

**Detailed materials and methods**

**Recruitment of animals for phenotyping in the experiment**

Careful selection of *desi* cattle was a critical aspect of this study to ensure accurate transcriptome profiling. Various exclusion criteria were applied to identify suitable cattle for inclusion, which involved excluding cows in advanced pregnancy or those that had received treatment with steroids or antibiotics within the last month. Additionally, cows suffering from any known chronic infections or debilitation were not considered. Cattles belonging to unwilling owners were also not included in the experiment.

For inclusion in the study, susceptible (diseased) and resistant (healthy) female cattle were meticulously chosen based on detailed criteria[[1–3]](https://paperpile.com/c/dMKByL/TNUM+sbSh+huDo):

*Inclusion criteria for susceptible (diseased) cows:* 1. History: Weakness, un-thrifty, diarrhoea (intermittent or continuous), reduced productivity. 2. Physical traits (a) Body Coat: Exhibited rough texture with dry skin, ribs visible from distance. (b) Body Condition: Animals showed signs of weakness and sustained weight loss. On a scale of 1-5, Body condition score (BCS) **<=**3. 3. Clinical symptoms (a) Weakness: weak cows (b). Diarrhea: cows suffering from chronic diarrhea not responding to treatment. (c) Feces: Loose 4. Production efficiency: (a) Lowered / reduced milk yield (b) Body weights: Static or decreased 5. Reproduction rate: increased age at first calving, long inter-calving period 6. Increased elimination from herd 7. Laboratory diagnosis (a) ZN Staining: Positive staining with categories +2 to +4.

*Inclusion criteria for resistant (healthy) cows*: 1. History: Healthy, no history of illness. 2. Physical Profile: cows in good health. 3. Co-morbidities: None 4. Clinical Symptoms & Diarrhea: Absence of clinical signs and no occurrences of diarrhea. 5. Productivity: Lactation average or above average, as per breed, stage of lactation and parity. (a) Body weight: Good and sustained body weights (b) Milk Production: Maintained (6) ZN Staining: Negative.

**Phenotyping of animals:**

Each cattle recruited for the project underwent phenotyping for Body Condition Score (BCS), clinical condition, and production parameters. Healthy animals were selected from the same farm, which had a prevalence of JD, and diseased cattle were recruited for the experiment. Healthy animals were phenotyped for over 18 months, and only those that remained healthy during this period were chosen as healthy or resistant animals for the transcriptome analysis in the project.

**Sample collection and RNA isolation**

Peripheral blood samples (5 ml each) were collected from phenotyped cattle, both those exhibiting the disease and healthy individuals, who met the inclusion and exclusion criteria outlined above. Samples were collected as per the guidelines of the Committee for the Purpose of Control and Supervision on Experiments on Animals (CPCSEA), India, with the Institutional Animal Ethics Committee (IAEC) approval. Following sample collection, peripheral blood mononuclear cells (PBMCs) were isolated using a density gradient centrifugation method with Hypaque-Ficoll (GE Healthcare Bio-Sciences AB, Uppsala, Sweden). Subsequently, PBMC samples were lysed using TRIzol reagent (Thermo Fisher Scientific, Waltham, MA, USA) and stored in a temperature-controlled chamber at −80°C. Total RNA extraction from purified PBMCs was carried out using the RNeasy Mini kit (Qiagen GmbH, Germany) according to the manufacturer's instructions. The integrity and quantity of the isolated RNA were assessed using a Bioanalyzer (Agilent Technologies, Inc).

**Library preparation and sequencing**

RNA libraries were prepared using the NEBNext Ultra RNA Library Prep Kit for Illumina (New England Biolabs Inc.), following the manufacturer's guidelines. Approximately 100 ng of RNA from each sample was employed for RNA library preparation. The quality of the resulting libraries was evaluated using a Bioanalyzer, and their quantification was performed using a Qubit 2.0 Fluorometer (Life Technologies) and quantitative PCR (qPCR). Subsequently, the libraries (1.3 ml each, 1.8 pM) underwent denaturation, dilution, and loading onto a flow cell for sequencing. The cDNA library preparation and Illumina sequencing were carried out on a Novaseq 6000 platform with 150 bp paired-end reads.

**Identification of lncRNA**

For lncRNA identification, a customized version of the LncEvo pipeline [[4]](https://paperpile.com/c/dMKByL/6plp) was employed, based on Nextflow. This pipeline was supplied with raw reads and an annotated Bos indicus reference genome (UOA_Brahman_1). The workflow initiated with data preprocessing using Fastp, followed by the removal of rRNA contaminations from each sample dataset. Subsequently, the cleaned reads underwent alignment to the reference genome using STAR. The resulting alignment files were then processed with Stringtie, generating GTF files containing assembled transcript structures for each sample. To refine the transcript dataset, all GTF files were merged using the Stringtie 'merge' function. Comparison with the reference GTF file was performed using CuffCompare [[5]](https://paperpile.com/c/dMKByL/HXrN) to eliminate transcripts with potential errors, denoted by class codes (c, e, p, s). Further stringent filtering steps were implemented: (i) Transcripts with class codes "=", "j," or "o" were discarded if the reference gene was not previously classified as lncRNA. (ii) Transcripts shorter than 200 bp in length were excluded. (iii) Strand-specific transcripts (-S option) and those with ORFs having a minimum protein length of 100 (-m 100), as identified by TransDecoder [[6]](https://paperpile.com/c/dMKByL/wpf5) , were also discarded. (iv) Transcripts classified as encoding by the Coding Potential Calculator (CPC) 2 [[7]](https://paperpile.com/c/dMKByL/SXvq) standalone version were removed from consideration. (v) Every transcript must exhibit an expression level meeting or exceeding 1 Transcripts Per Million (TPM) in at least one of the samples. Transcripts that passed these filtering criteria were identified as potential lncRNAs, resulting in the pipeline's output of a final GTF and FASTA file. This comprehensive approach ensures the reliable identification of long non-coding RNAs in the dataset.
